# Unique Asymmetric Branching of Drosophila Neurons Optimizes Temporal Dendritic Computation

**DOI:** 10.1101/2024.11.14.623502

**Authors:** Pin-Ju Chou, Ching-Che Charng, Harrison Hao-Yu Ku, Ann-Shyn Chiang, Chung-Chuan Lo

**Affiliations:** Institute of Systems Neuroscience, National Tsing Hua University; Department of Informatics, National Tsing Hua University; Brain Research Center, National Tsing Hua University

**Keywords:** Dendritic computation, asymmetric branching, structure-function relationship, order selectivity, coincidence detection

## Abstract

Neurons execute a versatile array of computations through a complex interplay of factors, including their morphology and synaptic architecture. Dendritic branching encodes upstream inputs into diverse spike patterns transmitted via downstream axons. While earlier studies highlighted the distinct morphologies and functions of a few representative neurons, the availability of large-scale electron microscopy and fluorescent imaging now enables comprehensive data analysis and further simulations to explore structure-function relationships more broadly. This study investigates the general morphological characteristics of diverse neuron types in the fly model. By employing the Strahler Order (SO) metric, we identified a specific bias towards asymmetry in neuronal branching and further investigated the effect of the asymmetry on computational capabilities. Specifically, symmetric branching enhances coincidence detection capability, whereas asymmetric branching increases input order-selectivity. While certain neurons exhibit extreme symmetry or asymmetry optimized for specific tasks, most neurons strike a balance between these computational strategies. This balance underscores the intricate relationship between neuronal structure and function. In contrast to the wide range of branching symmetries found in random bifurcation models, neurons across different species exhibit species-specific asymmetry, suggesting shared underlying branching mechanisms. Our findings provide a fresh perspective on the exploration of neuronal morphologies and their computational roles.

**Significance Statement:** This study reveals a novel structure-function relationship by analyzing the asymmetric branching patterns of fly neurons using extensive morphological data. While certain neurons display extreme symmetry or asymmetry for specialized computational roles, most converge toward a characteristic degree of asymmetry to balance their computational demands, especially in larger branching structures. This suggests that neuronal branching may be governed by intrinsic principles that support both developmental and functional needs.

## Main Text

### Introduction

Neurons exhibit various morphologies and perform versatile computational functions (1-3). Through the arrangement of synaptic decorations, primarily along dendritic branches, upstream signals are integrated and translated into fine-tuned spiking patterns (4, 5). Taking the intact branching structure into consideration, rather than a point neuron model, a single pyramidal cell can be likened to a multi-layered artificial neural network (2). Understanding the interplay between the branching structure and its computational functionality is a fundamental question in neuroscience.

Since the inception of neuroanatomy by Cajal, the diversity of neuronal shapes has become a key descriptor for neuronal classification, reflecting distinct functionalities and connectivity patterns (6-10). The formation of these various branching patterns necessitates a balance among several factors, including extrinsic neurotrophic gradients, intrinsic constraints related to wiring and transportation costs (11, 12), and the need to establish adequate connectivity for neural computations (13). Therefore, these complicated interactions seem to generate unique intricate branching patterns that lack clear similarities. In contrast to previous efforts that have focused primarily on a few representative neuron types, such as pyramidal cells in mammalian models (5, 14) and peripheral sensory dendritic arborization (da) neurons in the *Drosophila* larva model (15-26), the advent of large-scale collections of neuronal morphologies now enables the statistical discovery of general morphological features across a broader spectrum of neuron types (27-29).

Previous endeavors have underscored the crucial nexus between neuronal morphology and computational functions (30-34). Morphology alone significantly impacts spiking patterns, as demonstrated both *in vitro* and *in silico* (31-34). Compartmentalized dendritic branches enhance computational complexity in a branch-specific fashion (35, 36), shaped by the synaptic arrangements along these structures (35, 37). Conversely, physiological activity also promotes structural changes, such as those involved in learning (38). Thus, it is critical to investigate how branching patterns affect various computational scenarios for sensory information detection, including coincidence detection (33) and order selectivity (13, 39), which are crucial functions mediated by dendritic integration. This raises a key question: If neurons share general morphological features, how do these features relate to their computational functions?

In the study, utilizing the databases, FlyEM (29) and FlyCircuit (27, 28), each containing morphological data for over 20,000 fly neurons—approximately one-sixth of an entire fly connectome, we were allowed to exploit the comprehensive data investigating the underlying branching rules governing the neuronal morphology by statistical approach. Compared to artificially generated neurons, our results suggested fly neurons exhibit distinct congruent asymmetry, particularly in neurons with a higher number of branches. Cross-species observations reveal that neurons display a convergent yet species-specific degree of asymmetry.

To further our understanding of the structure-function relationship of this asymmetric property, we conducted computational simulations to explore the computational advantages associated with branching asymmetry. Our findings indicate that full symmetry enhances coincidence detection of upstream inputs, whereas asymmetric branching facilitates order-specific selectivity in strength. Furthermore, such specific branching asymmetry balances the trade-off between divergent computational scenarios.

## Results

### General Asymmetric Branching in the Shape of Fly Neurons

Neurons display intricate branching patterns to form connections dictating their computational potential (4, 40). To delineate the features underlying various neuronal morphologies, we harnessed big data from the FlyCircuit database (27) and the hemibrain dataset (29). Our primary focus was on the branching topology of fly neurons, as presented using the Strahler Ordering (SO) system (41, 42). Compared with the conventional Branching Ordering (BO) system (41), which determines branch order by the number of bifurcating nodes between the branch and the soma (Fig. S1A), the SO system assigns rank from the terminal nodes backward (Fig. S1B). In detail, terminal nodes are assigned SO1, and a branch’s SO increases by 1 recursively when its downstream branches are in the equivalent order; otherwise, it retains the highest downstream SO (Fig. S1B). Thus, the root (soma) has the highest SO, called the Strahler Number (SN). For neurons with equal branch numbers, a higher SN indicates greater symmetry (Figs. 1A and S1C).

**Figure 1.**
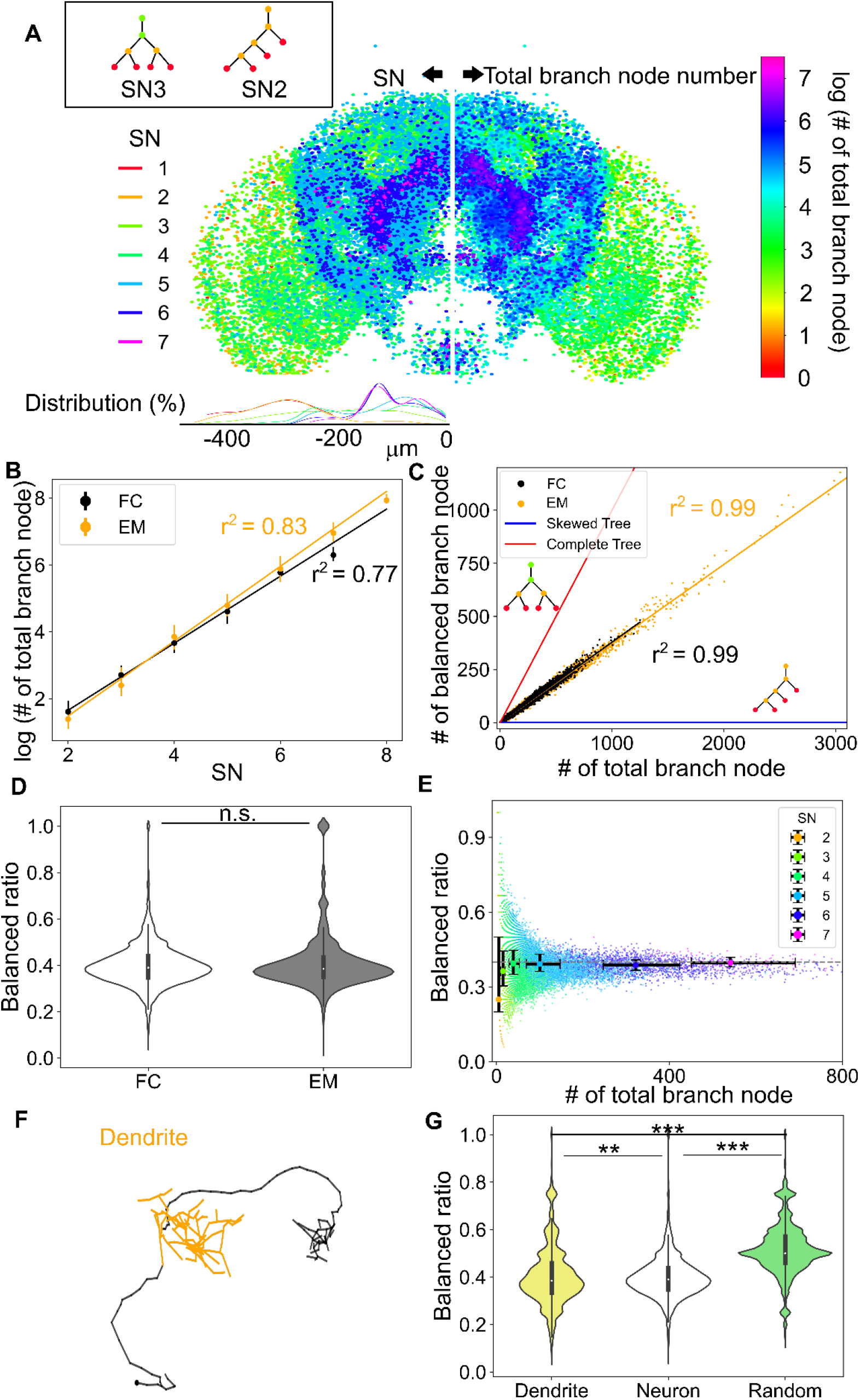
General asymmetric branching architecture of fly neurons. **A**, (Left) The somata of fly neurons with varying SN levels are unevenly distributed throughout the brain space. The inset shows neurons with the same number of branches but different symmetries (Left: a symmetric complete tree; Right: an asymmetric skewed tree). (Right) The somata distribution of fly neurons with varying branch numbers, shown on a log scale. Neurons on opposite sides are mirrored, with somas with SN projected from right to left and those with varying branch numbers from left to right; therefore, both sides of the brain contain all neuronal somata. **B**, The total number of branch nodes plotted on a log scale against SN levels, along with the fitted regression lines for both the EM (the hemibrain dataset (29), r^2^ = 0.83) and FC (the FlyCircuit database (27), r^2^ = 0.77) datasets. **C**, The number of branch nodes plotted against the number of balanced branch nodes, with fitted regression lines for the EM (r^2^ = 0.99), FC (r^2^ = 0.99), asymmetric skewed trees, and symmetric complete trees. **D**, Comparison of the balanced ratio distributions between EM and FC datasets (Mann-Whitney test, p > 0.05). **E**, The balanced ratio of each FC neuron plotted against the total number of branch nodes. Colors indicate different SN levels. **F**, A selected example of an FC neuron, with its dendrite (yellow) extracted using the NPIN algorithm (57). **G**, Comparison of the balanced ratio distributions among extracted dendrites, whole neurons, and a random growth model (see Methods). (Kruskal-Wallis test with post hoc Bonferroni test, *: p < 0.05, **: p < 0.01, ***: p < 0.001)

Using the Strahler Ordering metric, the SN distribution of fly neurons centers around mid-level SN ranks (SN4 and SN5) in both datasets, ranging from SN1 to SN7 in the FlyCircuit database and from SN2 to SN8 in the hemibrain dataset (Fig. S2, A and B). Surprisingly, neuronal somata with different Strahler Numbers (SNs) exhibit distinct spatial distributions (Figs. 1A, S2C, and S2D) — those with higher SNs are primarily located in central brain regions, while those with lower SNs are distributed more laterally. A higher SN indicates greater symmetry or a larger number of branches. We first investigated whether this distinct distribution suggests that neurons have varying numbers of branches in different brain regions. Astonishingly, we uncovered a mirrored distribution of the logarithm of branch numbers that closely resembles the distribution of SNs in both FlyCircuit (FC) (27) and the hemibrain datasets (EM) (29) (Figs. 1A and S2C). The branch number shows a strong correlation with SN (Fig. 1B, r^2^ = 0.77 for FlyCircuit data and r^2^ = 0.83 for the hemibrain dataset). Therefore, the SN of these fly neurons appears to be primarily influenced by the number of branches, with a similar degree of branching symmetry.

To further characterize the symmetry of neuronal branching, we defined each branching node as either balanced or imbalanced based on whether the SO level is upgraded relative to its downstream branches (Fig. S1B). We discovered a strong linear correlation between the number of total branch nodes and imbalanced branch nodes (Fig. 1C, r^2^ = 0.986 for FlyCircuit data and r^2^ = 0.992 for the hemibrain dataset), suggesting an underlying convergent branching asymmetry. By calculating the ratio of balanced branch nodes to the total number of branch nodes, we defined a metric called the ‘balanced ratio’. The theoretical upper bound of the balanced ratio corresponds to a complete tree topology, while the lower bound corresponds to the topology of a skewed tree. The balanced ratio of fly neurons centers at around 0.4 in both datasets (Fig. 1D), without significant differences (Mann-Whitney test, p > 0.9). Further analysis of the correlation between branch number and the balanced ratio reveals that neurons with a higher number of branches tend to converge more closely to the specific symmetry value, particularly starting from SN4 neurons (Fig. 1E, Kruskal Wallis Test, p < 0.001). On the other hand, to rule out the effect of non-bifurcating twigs, we trimmed the neuron by removing the non-bifurcating composition. The result suggests no effect of the existence of non-bifurcating branches (Fig. S3).

Next, to investigate the functional relevance, we analyzed the balanced ratio of neurons classified by neurotransmitter type, using GAL4 lines. The results show a significantly discrepant degree of asymmetry among neurons of different neurotransmitter types with mean values ranging from 0.36 to 0.4 (Fig. S3). Among these, npf-GAL4 neurons exhibited the highest symmetry, while Trh-GAL4 neurons, associated with serotonin, displayed greater asymmetry.

### Dendrites Possess Similar Asymmetricity with Whole Neurons

Dendritic branching structure plays a cardinal role in integrating upstream excitatory and inhibitory signals spatially and temporally (4). The synaptic decoration patterns along the dendritic twigs fine-tunes the resulting input signals (4, 13). Therefore, we took a further step to investigate whether neuronal dendrites, which are derived from its parental skeleton (see Methods, Fig. 1F), exhibit a similar degree of asymmetry. Our results suggest that the extracted dendritic branching structure retains a degree of asymmetry, significantly different but negligible from the original (Fig. 1G, Mann-Whitney test, p < 0.01. However, the effect size is negligible, |r| = 0.01). Notably, the asymmetry of dendrites of real fly neurons is significantly different from that of a randomly bifurcating construct with the same branch number (Fig. 1G, Mann-Whitney test, p < 0.001, the effect size is strong. |r| = 0.42).

### Boosted Coincidence Detectivity via Symmetric Branching Structures

Axonal spike generation in neurons arises from the dynamics of voltage-gated ion channels, particularly in the distal axonal initial segment (AIS) (43), driven by the spatiotemporal integration of synaptic inputs from activated synapses distributed along the dendritic branches. To investigate how asymmetric branching shapes dendritic computation and synaptic integration, we designed two key computational tasks: (i) coincidence detection, assessing the neuron’s response to simultaneous inputs, and (ii) order discrimination, evaluating the neuron’s ability to differentiate between stimuli presented in distinct temporal sequences.

To isolate the impact of asymmetric branching on neuronal coincidence detection, we first simplified the neuronal morphology by setting all branch lengths to equal values, making topology the only determining factor. Next, we injected currents simultaneously at each pair of two terminal points and quantified the coactivation strength relative to the sum of individual activations (Fig. 2A, see Methods). Our investigation of artificial tree structures revealed that skewed trees exhibited the lowest coincidence detection ratios, while symmetrical, fully balanced trees showed the highest. After normalizing the ratios based on the artificial trees with the same number of branches to eliminate the effect of neuron size, our results indicated a positive correlation between the coincidence detection score and the degree of branching symmetry (Fig. 2B, Pearson correlation: r = 0.547).

**Figure 2.**
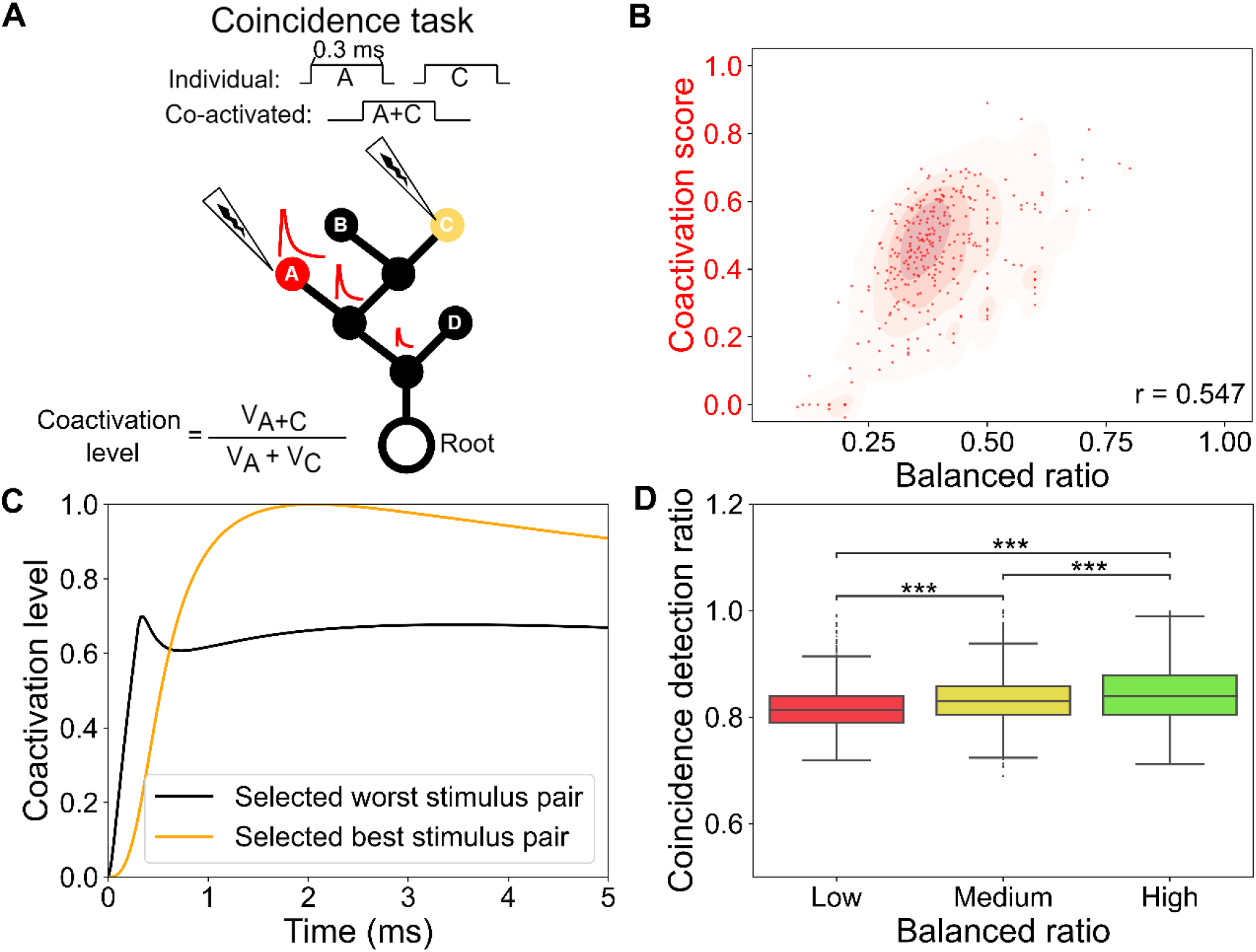
Symmetric branching structure enhances coincidence detectivity. **A**, Schematic representation of the coincidence task design. **B**, Coactivation scores from the topological simulation (branch length is neglected, and coactivation levels are normalized relative to the symmetric complete tree and the asymmetric skewed tree with the same number of branch nodes). Scatter points represent randomly sampled data for visualization. (Pearson Correlation, r = 0.547) **C**, Example traces of the best and worst coactivated pairs in a multi-compartmental simulation using the NEURON simulator (see Methods). **D**, Comparison of the coincidence detection ratios among low, medium, and high balanced ratio groups in a multi-compartmental simulation using the NEURON simulator (see Methods). (Kruskal-Wallis test with post hoc Bonferroni test, *: p < 0.05, **: p < 0.01, ***: p < 0.001)

To extend these findings to more realistic scenarios, we constructed multi-compartmental models of neuronal dendrites, preserving original branch lengths, using the NEURON simulator (Fig. 2D). In this case, since ideal artificial contrasts between skewed and symmetrical trees were unavailable, we based our analysis on neurons with similar effective sizes (see Methods). The selected example traces illustrating the coactivation levels are shown in Fig. 2C. The biophysical simulations consistently demonstrated that neurons with higher symmetry showed better coincidence detection (Fig. 2D). We also expanded the analysis to four activation points, further supporting the conclusion that symmetrical branching enhances coincidence detectivity.

### Augmented Order Selectivity with Asymmetrical Branching

Next, we investigated neuronal order selectivity by examining the ability of dendritic structures to distinguish between inputs with reversed temporal orders based on their resulting peak voltage changes (e.g., Stimulus 1 followed by Stimulus 2, and Stimulus 2 followed by Stimulus 1, Fig. 3A, see Methods). For each input pair within a structure, the number of distinguishable pairs was divided by the total number of pairs to calculate the order differentiation ratio (Fig. 3A). To highlight the impact of branching asymmetry, we initially equalized all branch lengths, focusing on the effects of branching asymmetry alone. Our analysis of artificial tree structures revealed that skewed trees exhibited the highest order differentiation ratios, while symmetrical trees had the lowest. The normalized order selectivity scores, ruling out the size effect by using artificial trees with the same number of branches, showed a negative correlation with the degree of branching symmetry (Fig. 3B, Pearson correlation: r = -0.704).

**Figure 3.**
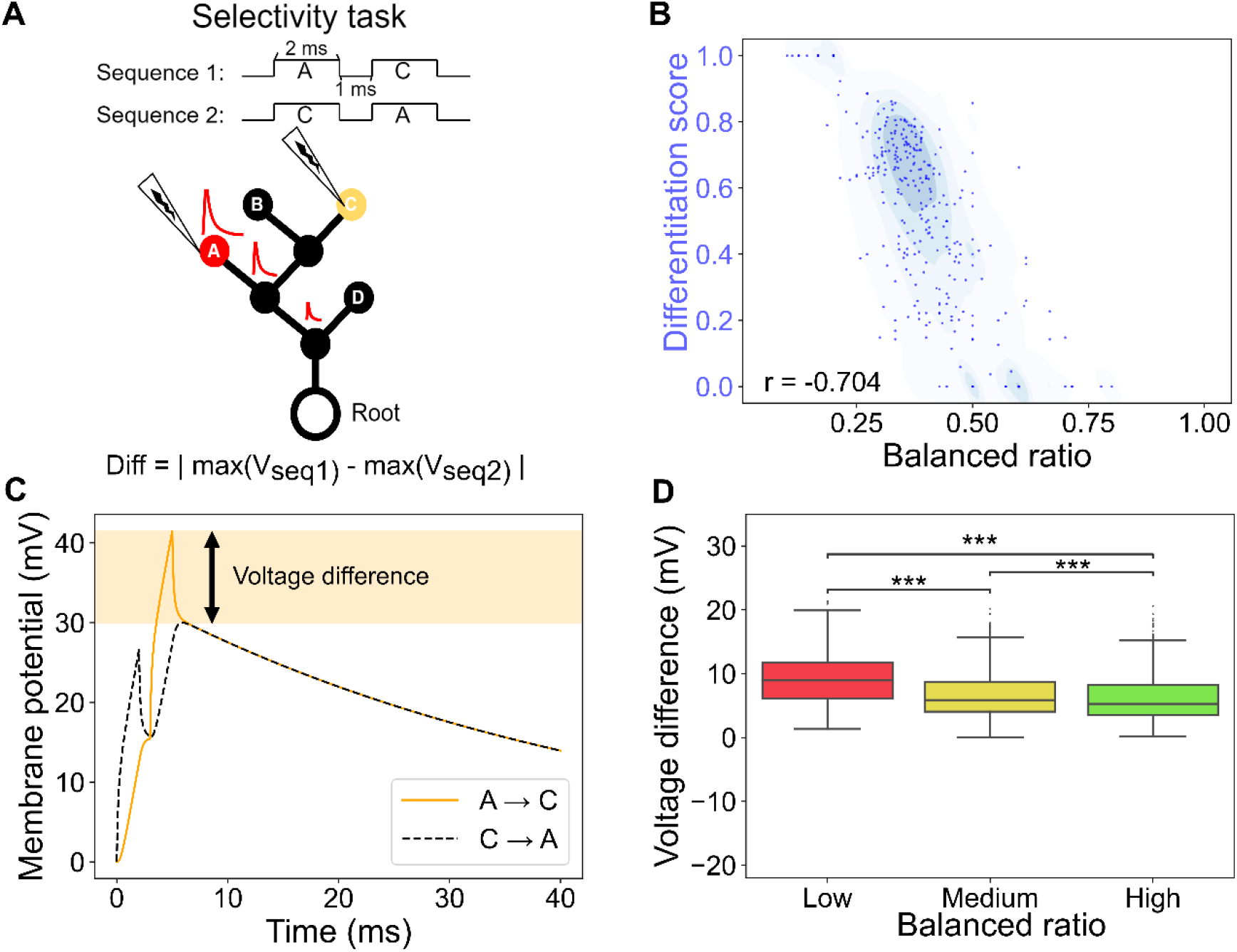
Asymmetric branching structure enhances order selectivity. **A**, Schematic representation of the selectivity task design. **B**, Differentiation scores from the topological simulation (branch length is neglected, and differences are normalized relative to the symmetric complete tree and the asymmetric skewed tree with the same number of branch nodes). Scatter points represent randomly sampled data for visualization. (Pearson Correlation, r = -0.704) **C**, Example traces of two reversed stimulation sequences in a multi-compartmental simulation using the NEURON simulator (see Methods). **D**, Comparison of the top 10% voltage differences among low, medium, and high balanced ratio groups in a multi-compartmental simulation using the NEURON simulator (see Methods). (Kruskal-Wallis test with post hoc Bonferroni test, *: p < 0.05, **: p < 0.01, ***: p < 0.001)

We extended this investigation by replicating the order differentiation task in NEURON-based multi-compartmental models that preserved the original branch lengths. In the absence of ideal artificial tree contrasts, we focused on neurons with similar effective sizes (see Methods). The selected example traces highlighting the voltage differences from the reversed stimulation sequence are presented in Fig. 3C. With the simulative model, we can quantify the difference between orders via voltage responses. The simulations confirmed that dendritic structures with lower symmetry exhibited larger voltage differences (Fig. 3D), indicating a stronger ability to distinguish between inputs arriving in different temporal sequences. This trend was further validated in scenarios with four sequentially arriving inputs, where neurons with more asymmetric branching consistently demonstrated enhanced order selectivity.

### Balance between Coincidence Detectivity and Order Selectivity

The degree of branching symmetry shows positive and negative correlation with the coincidence detection score and order differentiation score, respectively. By fitting the results using regression models, we obtained the convergent point which two lines meet (Fig. 4A). The balanced ratio of the intersection point is 0.42 which is close to the convergent asymmetry degree as we observed in larger neurons. In contrast, randomly connected branch structures with a balanced ratio mean of 0.52 showed a bias toward performance in coincidence detection (Fig. 4B). The significant difference between realistic neurons and randomly generated structures highlights the importance of unique asymmetry in neuronal branching patterns.

**Fig. 4.**
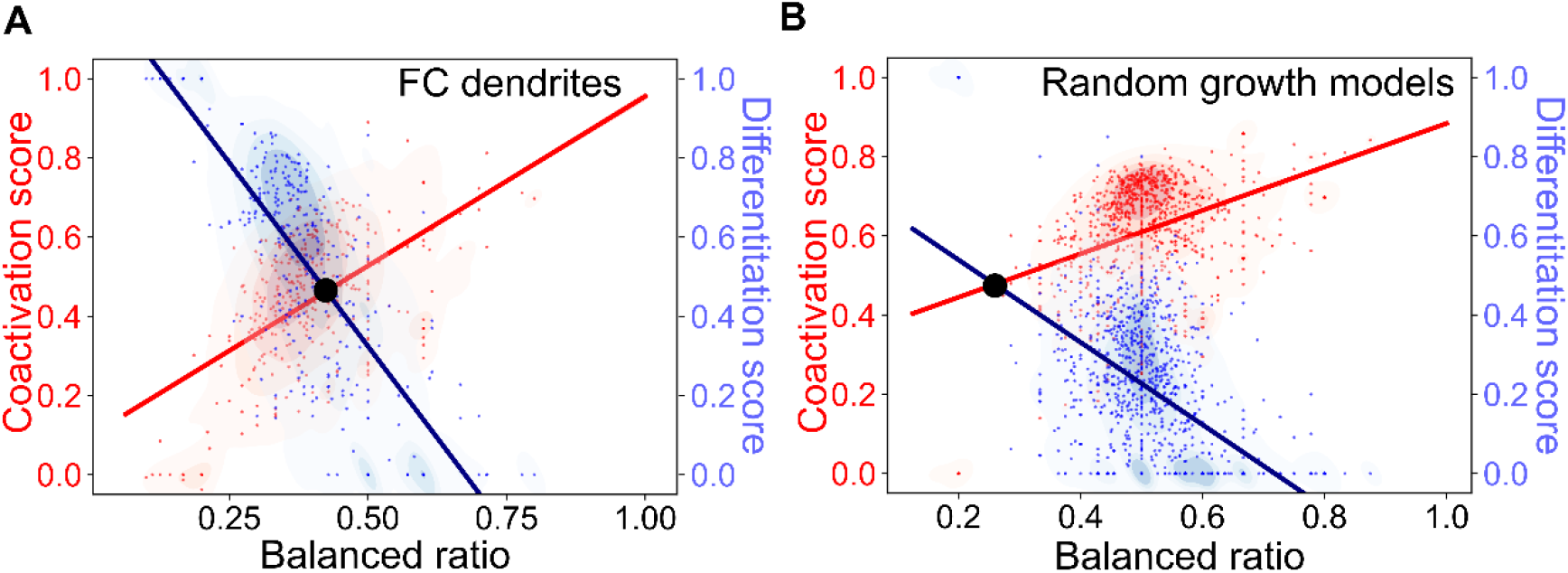
Tradeoff between coincidence detectivity and order selectivity. **A-B**, The distributions of coactivation scores and differentiation scores, as individually presented in Fig. 2 and Fig. 3, respectively, with fitted linear regression lines for FC dendrites (A) and random growth models (B). The balanced ratio at the intersection points is 0.42 for FC dendrites (A) and 0.26 for the random growth models (B). Scatter points represent randomly sampled data for visualization.

### Unique Asymmetry of Neuron Branching Structure

To examine the uniqueness of asymmetry observed in fly neurons, we constructed random bifurcation models, some of which have been proposed before (42, 44-46), based on renewability and parallelism for comparison (see Methods): Renewability determines whether a terminal node can repeatedly choose to bifurcate, transforming into a branch node, or has only one chance to either bifurcate or terminate permanently. Parallelism defines whether all terminal nodes choose to bifurcate simultaneously in each round or sequentially, one at a time (see Methods). Our results show that these models, across different uniform bifurcation probabilities, display a broad spectrum in the distribution of the balanced ratio (Fig. 5A).

**Figure 5.**
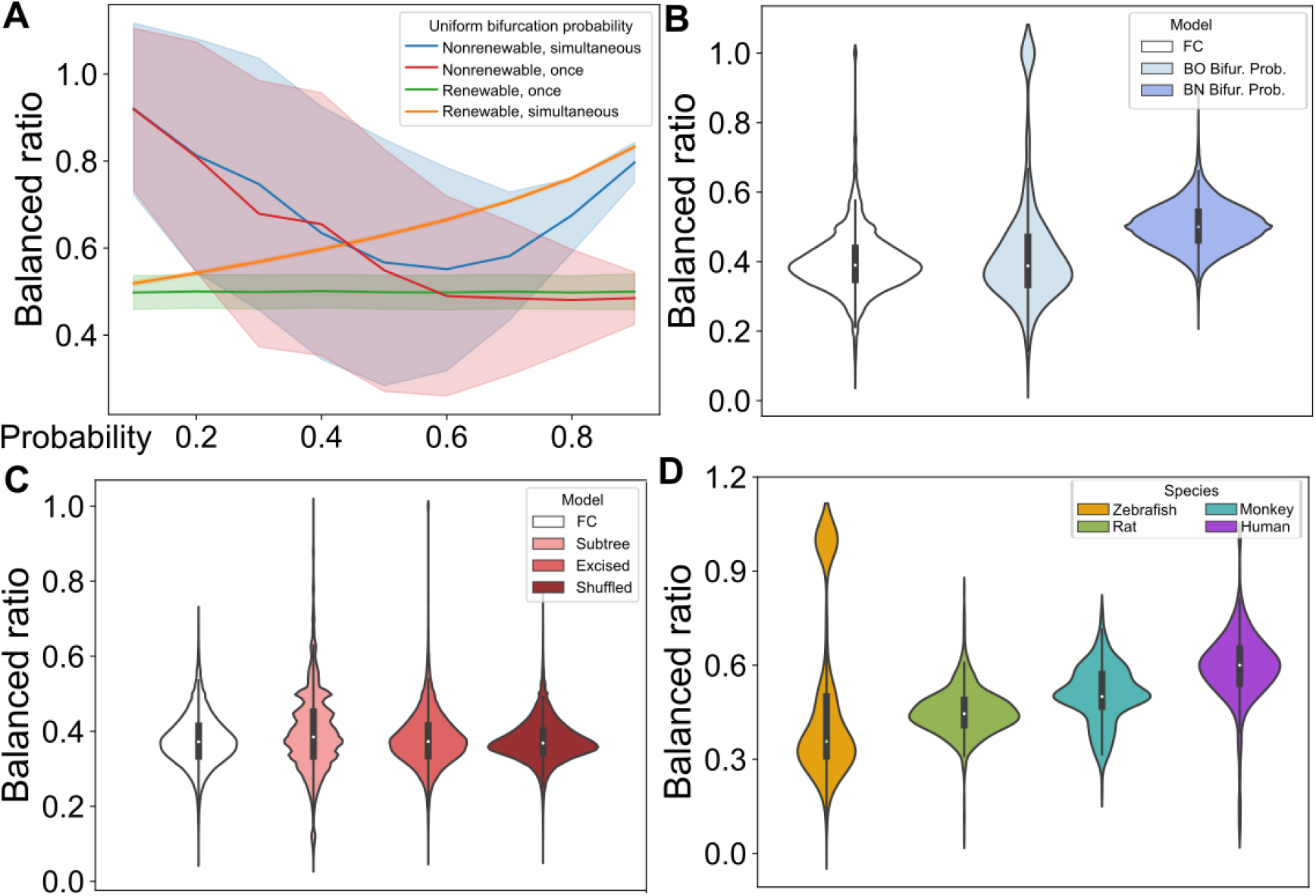
Unique balanced ratios of neurons across species rather than random bifurcation models. **A**, Balanced ratio distribution for four random bifurcation models, each generated with varying uniform bifurcation probabilities. The models simulate neuron branching based on principles of branching renewability and parallelism (see Methods). **B**, Balanced ratio distribution for random models informed by observed features of neurons from the FlyCircuit (FC) database. In the BO model, neurons bifurcate according to the bifurcation ratio at each branching order (BO) across different Strahler numbers (SN) (see Methods). The sampling ratio for each SN is derived from the original FC population. In the BN model, neurons bifurcate based on the observed bifurcation ratio at each branch node number (see Methods). **C**, Balanced ratio distribution for modification models, which are based on the original neuronal structure. In the subtree model, the extracted structure includes all downstream branches from a randomly selected node. In the excised model, the structure is obtained by severing a node from the original neuron. In the shuffled model, nodes and their downstream structures are randomly swapped with another node sharing the same branching order (BO) (see Methods). **D**, Comparison of balanced ratio distributions across different species, derived from the Neuromorpho database.

Therefore, we directly observed the bifurcation ratio along BO levels for a neuron and the branching probability according to the branch number. The bifurcation ratio along BO follows an exponential decay and stabilizes at a fixed ratio. By using the bifurcation ratio of neurons with different Strahler Numbers (SNs) as the bifurcation probability, we reproduced a balanced ratio distribution similar to that of fly neurons, with significant but negligible differences (Fig. 5B).

To further investigate whether the branching structure consists of homogeneous asymmetric components or a chimera of both asymmetric and symmetric elements, we developed subtree, excised, and shuffled models (see Methods). In the subtree model, we randomly extracted a subtree to assess the asymmetry within that section. The excised model involved randomly removing a skeleton node to observe its effect on branching asymmetry. In the shuffled model, we randomized the arrangement of downstream branches while preserving their original branch order (BO). Our results show significant but negligible populational differences, suggesting a more homogeneous configuration within the overall branching structure (Fig. 5C).

Then, a crucial question arises: Do neurons across different species also prefer a certain degree of branching asymmetry? To explore this, we extended our analysis to neurons from zebrafish, rats, humans, and monkeys using the Neuromorpho database (47). Our results suggest that the preference for a unique degree of asymmetry, though the value is different, is not limited to fly neurons but is shared across multiple species (Fig. 5D), indicating that neurons may follow similar fundamental branching mechanisms.

Next, we employed a metric, the balancing factor (bf), to evaluate the trade-off between wiring cost and transportation cost from the soma, assessing the intrinsic constraint (11) (see Methods). Our results suggest that the balanced ratio increases as more weight is given to the transportation cost (Fig. 6A-B). Consequently, wiring cost tends to be more prioritized in most neurons.

**Figure 6.**
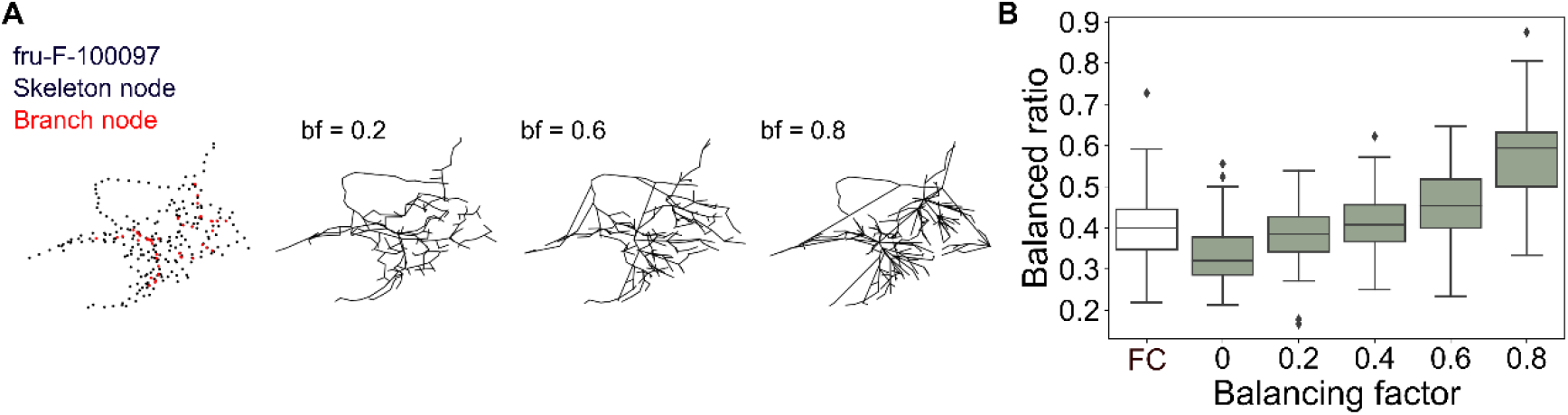
Trade-off between wiring and transportation costs. **A**, Example of a neuron and its reconstructions generated using different values of the balancing factor (see Methods). **B**, Comparison of balanced ratio distributions across varying balancing factor values.

## Discussion

Branching structure is key to bridging connectivity and computation. The rise of big data has enabled a statistical understanding of neuronal bifurcation principles. Our analysis of fly neurons, despite their diverse morphologies, reveals a convergent degree of asymmetry, particularly in neurons with more branches. Simulation tasks show that neurons balance order selectivity and coincidence detectivity, with asymmetric branching supporting the former and symmetric branching the latter, highlighting the significant impact of asymmetric branching on computation. In contrast to other random bifurcation models, only those following an exponential bifurcation propensity, as observed in neurons, can better reproduce this unique degree of asymmetry. Moreover, neurons from various species exhibit congruent asymmetry, suggesting shared fundamental branching mechanisms. In addition to branching asymmetry, most neurons balance the intrinsic costs of wiring and transportation at a specific ratio.

Coincidence detection and order selectivity are fundamental computational capabilities of neural computation, integrating synchronous and asynchronous signals, respectively. Simulated stimulation sequences show that symmetry alone impacts these two anti-correlated functions.

Coincidence detection is a general requirement for most neurons to overcome the firing threshold. On the other hand, order selectivity is a specialized function, as seen in the fly visual circuitry, where the asymmetric branching of T4 neurons, with a balanced ratio of 0.22—much lower than average— facilitates motion direction detection by enhancing order selectivity (48, 49). For most neurons, we found an intersection point where they achieve an optimized trade-off between these two tasks.

Morphological analysis often faces several challenges, including sampling bias in neuronal populations, limitations of imaging techniques, and tracing difficulties, among others. Surprisingly, despite discrepancies in branch number, fly neuron data from the FlyCircuit database and the hemibrain dataset display the same degree of branching asymmetry, cross-validating our findings. For neurons from other species in the Neuromorpho database, although the degree of asymmetry varies, they still adhere to a specific value, especially as branch number increases. Therefore, it is possible that neurons across different species share similar fundamental mechanisms and strategies for performing computations within neural networks.

Neuronal development is determined by the interplay of complex factors, including chemical gradients (50), adhesion proteins (51), intrinsic molecular network dynamics (18, 19, 21, 23, 24), and synaptic connectivity supported by surrounding partners (52). However, despite these intricate biochemical processes, neuronal morphology can be computationally reproduced by using observed morphological parameters (53, 54). Previous studies have primarily focused on typical neurons, such as pyramidal cells (42, 55) and Purkinje cells (42, 56) in mammalian models, and da neurons (15, 17, 26, 56) in the fly model. The branching architecture of da neurons has even been replicated through stochastic processes (15, 16, 23). Our study extends beyond these typical fly da neurons by statistically examining the branching topology of large datasets of neurons in the fly central brain. Our findings suggest that fly neurons exhibit a specific degree of asymmetry. The asymmetry distribution of these neurons can be reproduced through stochastic processes, with exponentially decaying branching probabilities converging at specific values, similar to those observed in pyramidal cells (55), while other commonly used random bifurcation models (45, 46) have failed in our simulation experiments. Considering the balance between intrinsic wiring costs and transportation costs (11), our results indicate that fly neurons have adopted a well-defined balance.

## Materials and Methods

### Strahler Order Calculation

In this system, each node is assigned a Strahler order (SO). Terminal nodes (nodes with no children) start with an order of 1 (SO1). For parent nodes, the order is determined iteratively: if all child nodes share the same SO, the parent’s order is one level higher; if child nodes have differing SOs, the parent inherits the higher SO from its child nodes. The highest SO in the structure, located at the root (soma), is designated as the Strahler number (SN).

### Balanced Ratio Calculation

Bifurcation nodes are classified as balanced or unbalanced based on the SOs of their child nodes; balanced nodes have child nodes with equal SOs. The balanced ratio is the count of balanced nodes divided by the total bifurcation node count.

### Dendrite Identification

The polarity of each skeleton node is identified by NPIN (57), a machine learning algorithm. Next, we used DBSCAN to further extract the dendritic arbor structures and validated manually.

### Balancing Factor for Evaluating Wiring and Transportation Costs

To construct optimized neuronal trees, the balancing factor which weighs wiring cost and transportation costs is applied (11). We used a minimum spanning tree algorithm to connect and incorporate the scattered skeleton points to the constructed tree structure. Each connection minimizes a total cost function that combines two elements: wiring cost (the Euclidean distance between each non-connected point and the closest tree node) and path length cost (the distance along the tree from the root to the skeleton point). The balancing factor (bf) adjusts the relative weight of these two costs, balancing material efficiency with conduction time. This single parameter directs the formation of the optimized tree structure.

## Acknowledgments

We appreciate the contribution of FlyCircuit and the hemibrain dataset. We also want to express our thanks to Dr. Li-An Chu and Dr. Chi-Hon Lee for their insightful comments.

